# Failure to account for behavioral variability significantly compromises accuracy in indirect population monitoring

**DOI:** 10.1101/2021.12.22.473935

**Authors:** Erin G. Wessling, Martin Surbeck

## Abstract

Indirect wildlife population surveying largely depends upon counts of artefacts of behavior (e.g., nests or dung). Likelihood to encounter these artefacts is derived from both artefact production and decay, and variability in production behavior is considered to contribute minimally to inaccuracy in wildlife estimation. Here, we demonstrate how ignoring behavioral variability contributes to significant population misestimation, using an example of an endangered ape, the bonobo (*Pan paniscus*). Until now, a single estimate of nest construction rate has been used to extrapolate bonobo densities, assumed to be broadly representative of bonobo sign production behavior. We estimated nest construction rates across seasons and social groups at the Kokolopori Bonobo Reserve, DRC, and find nest construction rates in bonobos to be highly variable across populations as well as seasonal. Failure to account for this variability led to degradation in the accuracy of bonobo population estimates of abundance, accounting for a likely overestimation of bonobo numbers by 34%, and at worst as high as 80%. With this example, we demonstrate that failure to account for inter- and intra-population behavioral variation compromises the ability to estimate both relative and absolute wildlife abundances. We argue that variation in sign production is but one of several potential ways that behavioral variability can affect conservation monitoring, should be measured across contexts whenever possible, and must be considered in population estimation confidence intervals. With increasing attention to behavioral variability as a potential tool for conservation, conservationists must also account for the impact that behavioral variability can play upon wildlife population estimation. Our results underline the importance of observational research to wildlife monitoring schemes as a critical component of conservation management. We discuss the avenues through which behavioral variability is likely to impact wildlife monitoring accuracy and precision and propose potential approaches for accounting for behavioral variability in wildlife monitoring.

## INTRODUCTION

Wildlife monitoring and assessments of population size are crucial components of biodiversity conservation. To effectively monitor species, the information gathered must be an accurate reflection of true status of a population while free of bias and precise enough to allow for differences in status to be informative (Kremen et al., 1994). Wildlife monitoring involves the quantification of direct or indirect observations of animals, which, in lieu of cost-prohibitive censusing, are commonly performed as surveying of subsets of the areas of interest. Sampling by direct observation has traditionally meant quantification of observations of animals by a surveyor (Buckland et al., 2001; Kühl, 2008), although technological and analytical improvements increasingly permit the use of remote methods to estimate animal abundances based on observations during camera trap or acoustic surveying (e.g., Campos-Candela et al., 2018; Cappelle et al., 2019; Crunchant et al., 2020; Howe et al., 2017; Moeller et al., 2018; Nakashima et al., 2018).

For especially elusive species or for surveying in dense vegetation, however, conservationists typically rely on the surveying of indirect signs of animal presence (Buckland et al., 2001; Plumptre, 2000), such as dung (e.g., Barnes, 2001; Marques et al., 2001; Massei & Genov 1998; Mayle et al., 1996; Nchanji & Plumptre, 2001; Plumptre, 2000; Rogers 1987), or remnants of behavior (e.g., nests; Kühl, 2008; footprints: Bonesi & Macdonald 2004). As in direct surveying, a great amount of attention centers around designing surveys to ensure sampling effort is sufficient and that animal counts are robust (Buckland et al., 2001). However, unlike direct surveying, the use of indirect surveying also necessitates accounting for auxiliary variables that account for potential sign abundance, such as rates of sign production and decay (Buckland et al., 2015). While the use of indirect surveying facilitates the possibility of surveying elusive wildlife populations, the additional consideration of sign production and decay represent a significant potential source of error, leading to inherent flaws in what still remains a fundamentally important methodology (Bailey & Putnam 1981, Hayward & Marlowe 2014). Some authors argue that this component of population estimation requires greater amounts of attention, as this is where potential biases are most easily introduced (Bailey & Putnam 1981; Strindberg et al., 2018).

For example, conditions of the local environment are a commonly acknowledged influence on the probability of sign encounter, and heterogeneity is common in metrics of sign decay rates across locations (e.g., Bessone et al., 2021; Kühl et al., 2007; Walsh & White, 2005). Sign decay has been linked to a number of variables such as climatic seasonality, construction material or dung matrix, storm frequency, and sun exposure (e.g., Bessone et al., 2021; Kamgang et al., 2020; Kouakou et al., 2009; Laing et al., 2003; Morgan et al., 2016; Nchanji & Plumptre, 2001; Plumptre, 2000). Therefore, it is commonly recommended that decay rates are measured locally during surveying, as failure to do so may result in imprecise measurement and hinder the validity of inter-site comparisons (e.g., Bessone et al., 2021; Kühl, 2008; Laing et al., 2003; Mohneke & Fruth, 2008).

However, an often-overlooked component of wildlife monitoring relates to the variation in the production of indirect signs, which is a derivative of behavior by the species surveyed. Rates of production behavior for many indirect signs are typically treated as static entities – derived from a single group (e.g., Hedges et al., 2005; Kouakou et al., 2009; Morgan et al., 2006; Todd et al., 2008), or even one or two individuals (Mitchell et al., 1985; Viquerat et al., 2012) — and considered representative for the species across multiple localities. Single measures are commonly considered sufficient because measurement of production behavior must be directly observed to be quantifiable, which is both frequently unfeasible during surveying while also negates the need for indirect surveying in the first place (as population size is observable and therefore known already to the surveyor). As indirect surveying rarely occurs when behavior of the population is directly observable, sign production behavior must typically be measured separately from the surveyed populations.

Nevertheless, given that behavior is frequently variable within a species it may be problematic to rely on a single measure to represent species-level patterns. Variation in animal behavior may be influenced by the environment (e.g., Andersen et al., 1992; Mitchell et al., 1985; Kalan et al., 2020) but can also vary without clear environmental drivers (e.g., Samuni et al., 2020), and is most frequently tied to seasonality (e.g., Mitchell et al., 1985; Mayle et al., 1996; Rogers et al., 1987; Todd et al., 2008). The scale of variability in sign production behavior is argued to be small and therefore evaluation of sign production variability is scant relative to drivers of sign decay variability (Marques et al., 2001; although see Todd et al., 2008). In few cases when variation in sign production is described, the impact of sign production variability remains relatively unevaluated in the context of its impacts on species population estimates. Consequently, what are the impacts of ignoring behavioral variability on the accuracy of absolute and relative wildlife population monitoring estimates?

To investigate the impact of behavioral variation upon issues of accuracy and precision in species monitoring, highly behaviorally flexible clades like great apes serve as ideal models. Apes are among the most extensively documented clades to exhibit behavioral variation and likely also among the most flexible (e.g., Kalan et al., 2020). As ape surveying has historically been conducted predominately via indirect surveying with little current methodological alternative (although emerging camera-trap methodologies increasingly permit monitoring, albeit on smaller scales; e.g., Bessone et al. 2021; Campos-Candela et al., 2018; Cappelle et al., 2019; Crunchant et al., 2020; Howe et al., 2017; Moeller et al., 2018; Nakashima et al., 2018), evaluating sign production variability in a species like the bonobo (Pan paniscus) represents a straightforward approach to understanding the impacts of extensive expressions of behavioral variability upon accurate population estimation. Indeed, in the case of ape nests, sign construction is known to vary according to weather patterns (Stewart et al., 2018), therefore it is already likely that we have ignored potential patterns of behavioral variation which affect ape density estimations. Bonobos are endemic only to the Democratic Republic of the Congo (DRC), and as one of the ape species under greatest threat, accurate population monitoring is critical at both the absolute and relative scales (Fruth et al., 2016). There are only an estimated minimum of 15-20,000 remaining individuals in the wild (IUCN & ICCN, 2012), although surveying is infrequent due to high logistical obstacles and therefore we know comparatively little about their current distribution. Consequently, large-scale models of bonobo abundance rely heavily upon few estimates of local densities. Under-surveying of bonobo populations has also led to an inability to reclassify the species as critically endangered (Fruth et al., 2016), meaning that accurate population monitoring is both of pressing need and a current conservation hurdle because data are scarce.

Bonobos, like all apes, construct nests to sleep in at night (Fruth & Hohmann, 1993), which is the predominant target of observation in bonobo surveys (Kühl, 2008). Bonobos regularly also construct nests for lounging during the day (Fruth & Hohmann, 1993), thereby providing ample opportunity for construction behavior to vary. While nest decay rates for bonobos have been measured at a few sites (Figure S1, Table S1), nest construction rate has to date only been measured in a single location (LuiKotale: Mohneke & Fruth, 2008). Furthermore, a portion of bonobo densities have also been estimated under the assumption of a single nest constructed per day (e.g., Hashimoto & Furuichi, 2002; Inogwabini et al., 2008; Reinartz et al., 2006; Van Krunkelsven, 2001). Meanwhile in chimpanzees, the sister species of bonobos, nest construction rates vary by ca. 5% across populations (Kouakou et al., 2009; Table A1). Generally, bonobos are argued to be comparatively less variable in their behavior than chimpanzees (Hohmann & Fruth, 2003) and occupy a considerably smaller and less environmentally variable biogeographic range (Fruth et al., 2016). Therefore, it may be expected that behavioral variation in nest construction is comparatively lower in bonobos than in chimpanzees. In this study, we first aimed to evaluate variation in nest construction rates, and secondly to evaluate the impact of behavioral variation of this trait on our ability to accurately estimate bonobo populations from nest counts. Specifically, we consider cross-site as well as intra-site (e.g., season, sex, social group) variation in nest construction behaviors, and re-evaluate published estimates of bonobo densities to account for the likelihood that both patterns of nest construction and decay can be variable.

## METHODS

To evaluate potential variability in nest construction behavior in bonobos, we collected data on the nesting behavior of three distinct social groups at the Kokolopori Bonobo Reserve in the DRC (Surbeck et al., 2017). We collected data during 410 full-day focal follows over the course of one calendar year (September 2020 – August 2021) on a total of 33 adult individuals (10 male, 23 female; mean days/individual:12.4 days, range: 3-25) from the three neighboring communities (Ekalakala, Kokoalongo, and Fekako) with a mean of 137 observation days per group (range: 77 – 172). During focal follows, observers marked each instance of nest construction and the species used to construct the nest. As observation was occasionally interrupted or focal animals were lost over the course of the day, we restricted all subsequent analyses to follows at least six hours in length that spanned the entirety of daylight hours (from morning nest to night nest) to reduce the likelihood that observations of nest construction were missed. The restriction of data that meet these criteria, therefore, reduced the dataset from 410 to 386 follow days.

Some researchers have previously argued that day nests are of flimsier construction than night nests and therefore should not be considered in calculations of nest construction rates (e.g., Fruth & Hohmann, 1993; Van Krunkelsven, 2001), however, most studies have nonetheless included day nests in the calculation of nest construction rate (e.g., Kouakou et al., 2009; Mohneke & Fruth, 2008; Morgan et al., 2006). Regardless of structural robustness at construction, because day nests still require the bending of branches in a manner that is indistinguishable from a night nest during surveying, we argue that they must be included in nest construction rate, as robustness of nest construction only relates to the durability (i.e., rate of decay) of the nest but not its identifiability. In this sense, future studies should measure if day nest durability differs from that of night nests.

To calculate average nest construction rate at Kokolopori, we fitted a Generalized Linear Mixed Model (GLMM; Baayen, 2008) with Poisson error structure, with the number of nests constructed during the course of a follow as the response. We tested a potential seasonal effect in nest construction behavior using the sine and cosine of the radian of Julian date of the focal follow (Stolwijk et al., 1999), as well as for sex differences in nest construction behavior by including the sex of the focal individual as a predictor. We accounted for potential group differences in nest construction behavior by including social group as a predictor (fixed effect) and included focal individual and date of the focal follow as random effects. To account for varying observational effort, we included the log of the duration of a focal observation as an offset term. We found no issues with model overdispersion (dispersion parameter = 0.42), collinearity among predictors, or model stability. We used the intercept of the model to derive an average nest construction rate for the population while correcting for all significant categorical predictors, if relevant. We compared the fit of the model to a null model lacking the test predictors of sex and seasonal terms (but otherwise identical) using a likelihood ratio test (Dobson & Barnett, 2018). We evaluated predictor significance similarly, by excluding each predictor and comparing each reduced model to the full model using a likelihood ratio test (ibid.). We assessed model stability by excluding each level of the random effects one at a time and comparing the estimates with those derived for the full data set. Lastly, we derived confidence intervals by means of parametric bootstraps (function bootMer of the package ‘lme4’, version 1.1.27.1; Bates et al., 2015).

Nest construction rates in bonobos have been previously described at only one site (LuiKotale; Mohneke & Fruth 2008), however these authors used a different calculation than that used here. Therefore, to contextualize our results in the context of other published nest construction rates, we also sought to verify that potential inter-site differences in nest construction rates could be attributed only to differences in behavior and not to methodological differences in rate calculation. Therefore, we also calculated average nest construction rate using Mohneke & Fruth’s (2008) calculation, which presumes sex differences in construction behavior and estimates an average construction rate based on average party sex ratios, using the party composition from group follows for the same period, using 293 days and 10635 30-min party composition scans.

If nesting behavior varies seasonally, surveying conducted during one period of an annual cycle may identify a greater number of nests than a survey conducted during another period of the year, despite no change in the number of nest constructors. Further, the time it takes for a sign to decay represents also the time window within which sign production behavior is relevant to each survey. Consequently, the nest decay period chosen as well as the date a survey was conducted may impact inter-survey comparability if nest construction behavior is temporally variable. Therefore, to better understand the seasonal variability in average nest construction behavior ultimately relevant for bonobo population monitoring, we used each of the four unique nest decay rates previously published for bonobos (Table A1) as a sampling window prior to each potential survey day during the year (n=365). Because we do not have multiple years of data, we treated date cyclically when sampling, e.g., using a 183-day decay rate the nest construction rate estimated on January 1 calculates a nest construction rate using data collected during focal follows on the last 183 days of the same calendar year). Then, we calculated the average nest construction rate for each combination of decay rate and date in the year.

Lastly, to contextualize the impact of variable nest construction rates on estimates of bonobo densities across their range, we considered the variability of published nest decay rates and nest construction rates for bonobos (Table S1) for their impact on published bonobo density estimations. Commonly, ape density estimates are derived using the following generalized equation: *D=N* / (*A* × *p* × *r* × *t*), where *N* is the count of nests discovered, A is the area surveyed, p is the proportion of nest builders within the population, *r* is the nest production rate, and *t* is the nest decay rate (Buckland et al., 2001; Kühl, 2008). We replaced original nest production and decay rates with all combinations of these values (including rates from Kokolopori derived here) and permuted all possible outcomes of density for each published non-zero density estimate. We additionally considered the effect of seasonal variation on nest construction rate in these permutations by allowing for density estimations to derive from a balanced sampling of either the single published value from LuiKotale (Mohneke & Fruth, 2008) or any of the possible seasonally variable construction rate values from our Kokolopori dataset, based on the decay rate of 76 days (Mohneke & Fruth, 2008). In other words, we allowed for a total of 730 possible nest construction values to be permuted within the model, the 1.37 nest/day from LuiKotale (n=365) and the seasonally varying nest construction rates from Kokolopori (n=365). ‘Thus, with this analysis we are evaluating the relative change in published density estimates when using different values of nest construction and decay.

## RESULTS

In our evaluation of Kokolopori nest construction rates and the factors that influence them, seasonality, group, and sex significantly contributed to explaining variation in nest construction rate (full-null model comparison: χ2 = 26.28, df = 3, p < 0.001). Specifically, nest construction behavior at Kokolopori varied seasonally (χ2 = 24.31, df = 2, p < 0.001), with highest rates of nest construction during October (the wettest month of the year: Samuni et al., 2020) and the lowest number of nests produced during April. We did not find significant group or sex differences in nest construction rates (Table 1). As rainfall is a common predictor of variability in nest decay (Bessone et al. 2021), we also fitted an ad hoc model identical to our Poisson model, but replaced the generic seasonal predictor (sine and cosine of Julian date) with cumulative rainfall in the 4 weeks prior to each focal follow day. We found that rainfall significantly predicted variation in nest construction rate (full-null model comparison: χ2 = 5.179, df = 1, p=0.023; Table A2), with bonobos constructing more nests during periods of high rainfall. This pattern corresponded to a difference of 0.65 nests/day (range: 1.54 – 2.19 nests/day) over the range of monthly rainfall patterns at the site (range: 7-221mm cumulative rainfall).

**Table 1.** Effect of season (represented by sine and cosine of the radian of Julian date), sex, and group on nest construction behavior of two social groups of bonobos at the Kokolopori Bonobo Reserve in the Democratic Republic of the Congo (n = 268; 12 mo., 2020-2021), using a GLMM model with Poisson error distribution.

| Predictor | Estimate ± SE | CI<sub>(95%)</sub> | $\chi^2$ | p-value |
| --- | --- | --- | --- | --- |
| Intercept | -1.63 ± 0.06 | -1.761, -1.525 | - | - |
| Sine | -0.26 ± 0.05 | -0.368, -0.147 | -4.869 | <0.001 |
| Cosine | 0.05 ± 0.05 | -0.063, 0.154 | | |
| Group (Fekako)<sup>a</sup> | 0.13 ± 0.11 | -0.081, 0.328 | 1.494 | 0.473 |
| Group (Kokoalongo)<sup>a</sup> | 0.04 ± 0.08 | -0.115, 0.199 | | |
| Sex (Male)<sup>b</sup> | -0.11 ± 0.08 | -0.276, 0.053 | 1.670 | 0.196 |

<sup>a</sup>Reference category: Ekalakala; <sup>b</sup>Reference category: Female

When considering average observation duration (9.85 hours), we estimated that average nest construction at Kokolopori was 1.92 ± 0.06 nests per day (SE; model intercept, back-transformed), considerably higher than the previously published estimate from LuiKotale (1.37 nests/day: Mohneke & Fruth, 2008), as well as from the commonly assumed rate of 1 nest/day (Hashimoto & Furuichi, 2002; Inogwabini et al., 2008; Reinartz et al., 2006; Van Krunkelsven, 2001). When calculated following Mohneke & Fruth’s (2008) method, nest construction rate (1.92 nests per day) did not differ from the rate derived from the GLMM, indicating that differences in nest construction rate between published construction rate estimates from different research sites (i.e., Kokolopori versus LuiKotale) are not a methodological byproduct.

In our evaluation of the impacts of date and decay rate used in estimations of nest construction rates, we found that both impacted nest construction rates. Across the four nest decay sampling windows used, intra-annual variation in construction rate at Kokolopori averaged 1.04 nests/day over the year (range: 0.66 – 1.24 nest/day intra-annual range in construction rate), although average construction rate between estimates using each of the four unique nest decay rates varied minimally across different nest decay rates used (0.003 maximum difference between averages). All estimated construction rates at Kokolopori averaged higher than the previously published construction rate from LuiKotale (Mohneke & Fruth, 2008; Figure 2).

**Figure 1.**
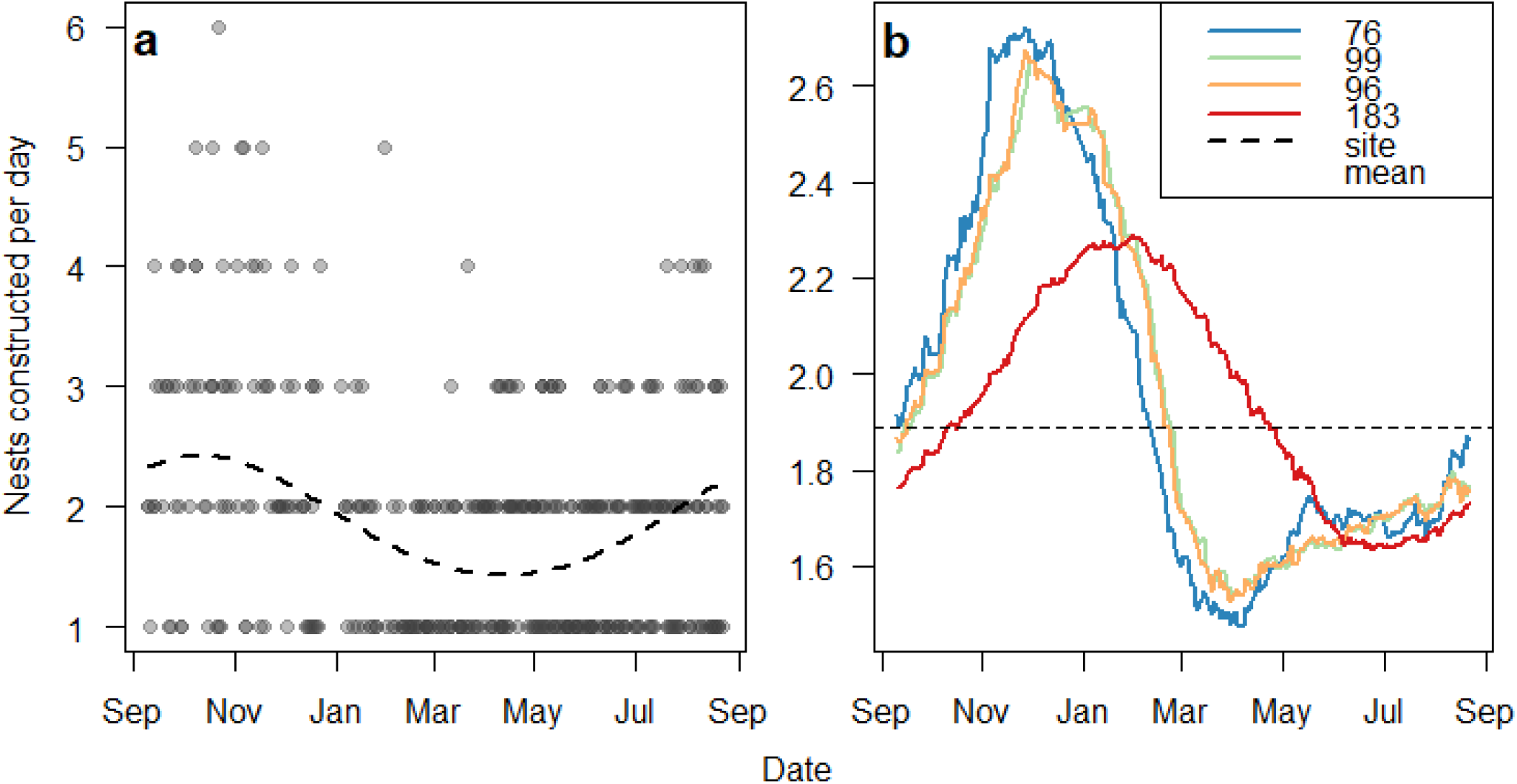
(a) Seasonal variation (represented by sine and cosine of radian of Julian date) of nest construction behavior in two bonobo communities at the Kokolopori Bonobo Reserve in the Democratic Republic of the Congo (12 months; 2020-2021). Circles represent the number of nests constructed by a single individual from dawn to dusk (focal follow), and the dashed line represents the model prediction derived from a GLMM model with Poisson error distribution. (b) Average Kokolopori nest construction rates estimated across four sampling windows (color-coded according to common nest decay rates [in days]).

**Figure 2.**
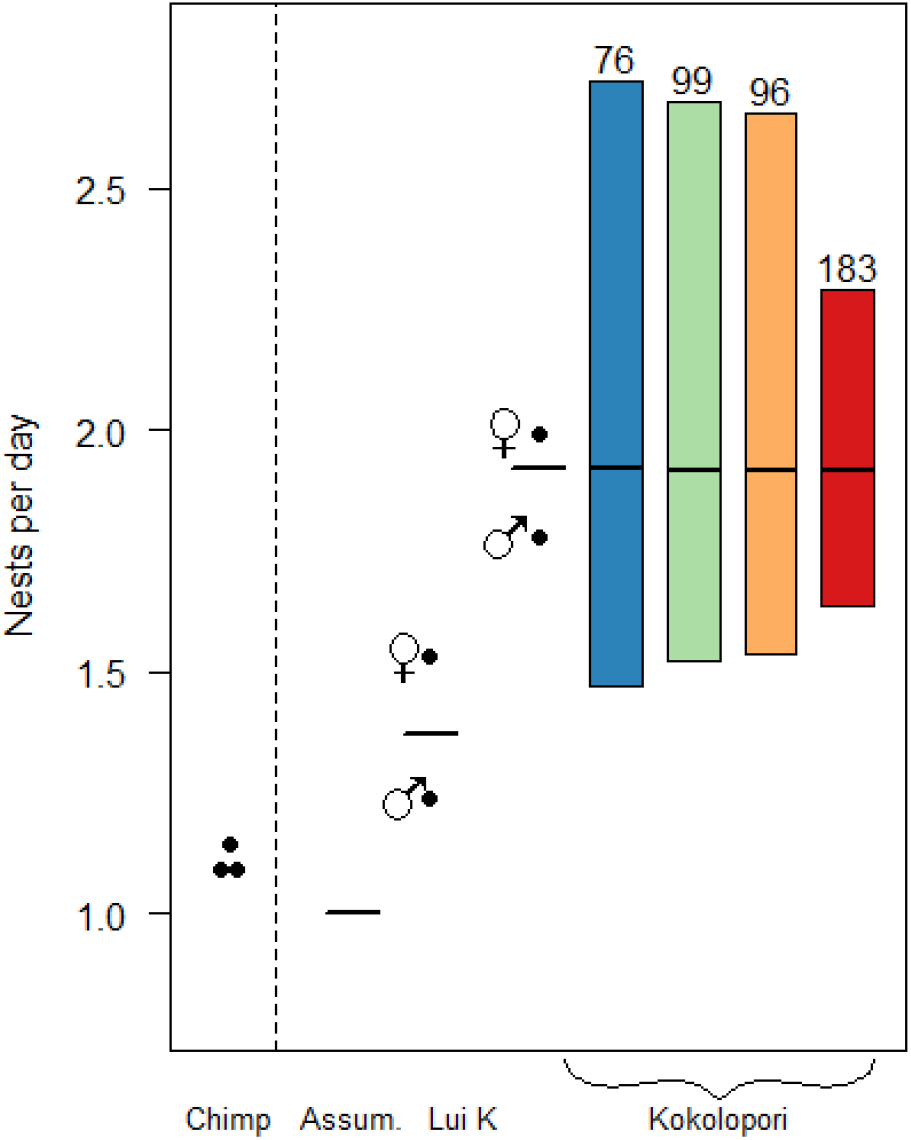
Nest construction rates from chimpanzees (‘Chimp’, left dots; Kouakou et al., 2009, Morgan et al., 2007, Plumptre & Reynolds 1997, various locations) and bonobos (right of vertical dotted line, various locations in Democratic Republic of the Congo). Bonobo construction rates include the common assumption of 1-night nest/day (‘Assum.’, left bar), calculated rates from LuiKotale males, females, and average (‘Lui K’, dots and bar; Mohneke & Fruth 2008), construction rate from bonobos at the Kokolopori Bonobo Reserve using Mohneke & Fruth’s (2008) calculation (dots and bar), as well as Kokolopori bonobo nest construction rates based on seasonal sampling using four nest decay windows (colored boxes, labeled by days to decay; 12 mo. study duration, 2020-2021.

To translate the impacts of behavioral variability to species-level monitoring, we evaluated the impact of different nest construction and decay rates on published bonobo densities across their range. In our analyses, we consider there to be potential bias (e.g., over- or under-estimation) in bonobo densities when we have identified disparities between published and permuted bonobo densities. Permuted bonobo densities across all potential bonobo nest construction rates were unanimously lower than originally published estimates, suggesting potential overestimation of densities in original values (Table 2). Potential overestimation of bonobo densities averaged 33 ± 5% (SD) (i.e., positive density bias) when permuted across all site-averaged construction rates, in the most severe case reaching up to an 80% positive bias. Additionally accounting for intra-annual variation in nest construction rates in our permutations reduced potential positive biases of rates minimally (1%) but increased the potential severity of positive bias in density estimates by up to 15%, as potential permuted nest construction values became increasingly variable and more seasonally extreme. When permuting densities across construction rates only, the five highest bonobo densities (Figure 3) suffered the highest positive bias (mean ± SD: 36.5 ±11%, range: 27 – 63%). Original densities were only potentially underestimated (% change) in cases where nest decay rates were permuted at a shorter decay rate than was used in the original study, specifically the density estimates from Serckx et al. (2014) which used the longest decay rate in our dataset of 183 days (Figure 3). Disparity between permuted and published density estimates became more severe when we permitted variation in both nest construction and decay, regardless of the direction of the misestimation.

**Table 2.** Average positive (underestimate) and negative (overestimate) changes in density estimates from originally reported values (Table S3) based on permutations of all potential nest decay and/or nest construction rates (Table S1), calculated either using site-wide averages or allowing for seasonal variation. Over- and underestimate changes were estimated separately. Note, a negative change in an estimated density would indicate that the original value was overestimated; values represent a change in permuted density relative to original density.

| Permutation type | [Overestimate]<br>Mean ± SE (%) | [Underestimate]<br>Mean ± SE (%) | SD<br>(%) | Range min, max<br>(%) |
| --- | --- | --- | --- | --- |
| Decay and construction (site average) | -38.2 ± 1.5 | 55.8 ± 9.4 | 38.3 | -71.7, 140.8 |
| Decay and construction (with seasonality) | -37.1 ± 0.1 | 60.0 ± 0.6 | 39.3 | -80.1, 140.8 |
| Construction only (site average) | -29.1 ± 2.8 | - | 16.4 | -47.6, 0.0 |
| Construction only (with seasonality) | -28.2 ± 0.2 | - | 17.4 | -63.3, 0.0 |
| Decay only | -33.5 ± 3.1 | 35.0 ± 4.4 | 40.6 | -58.5, 140.8 |

**Table 3.**
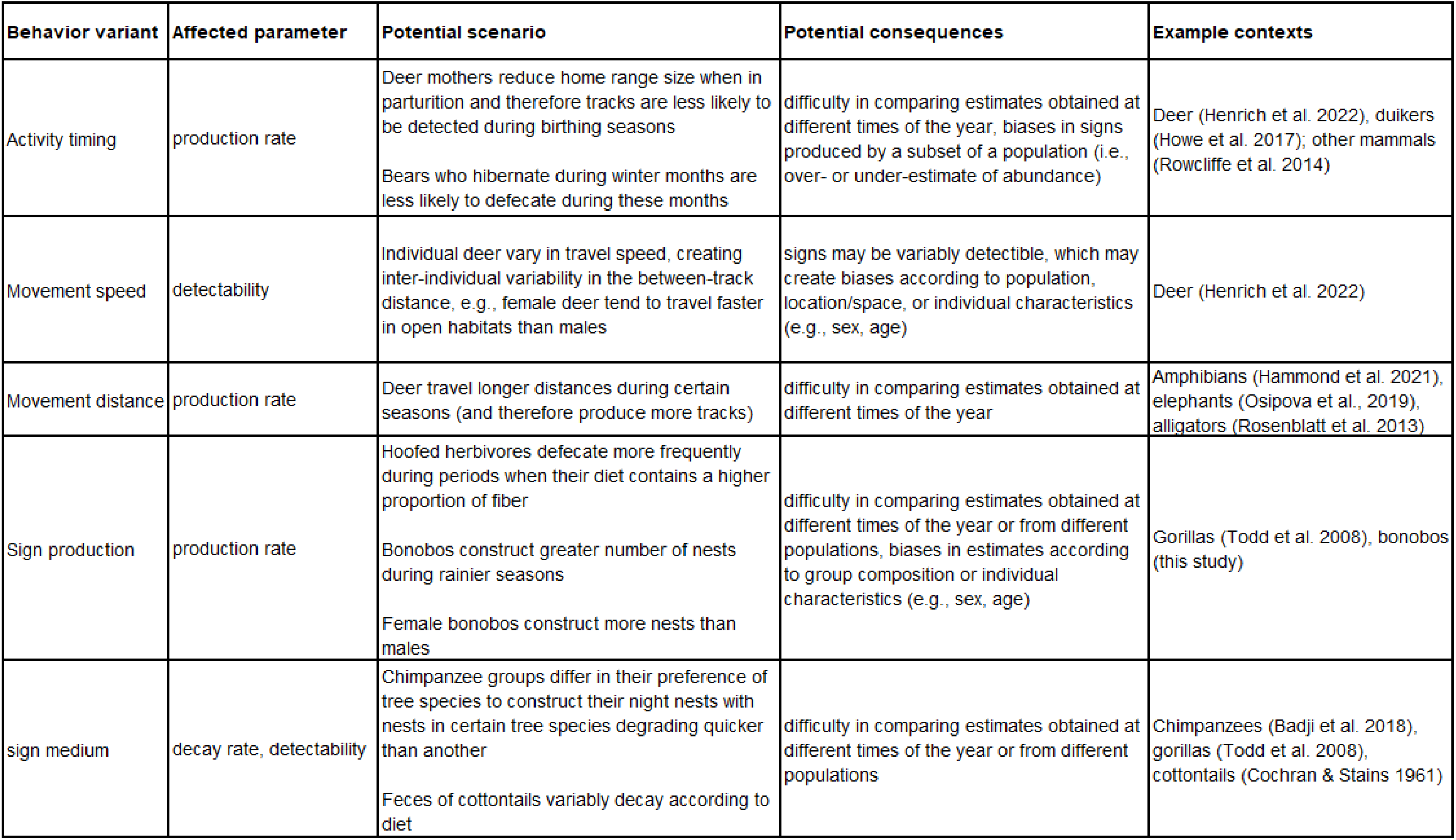
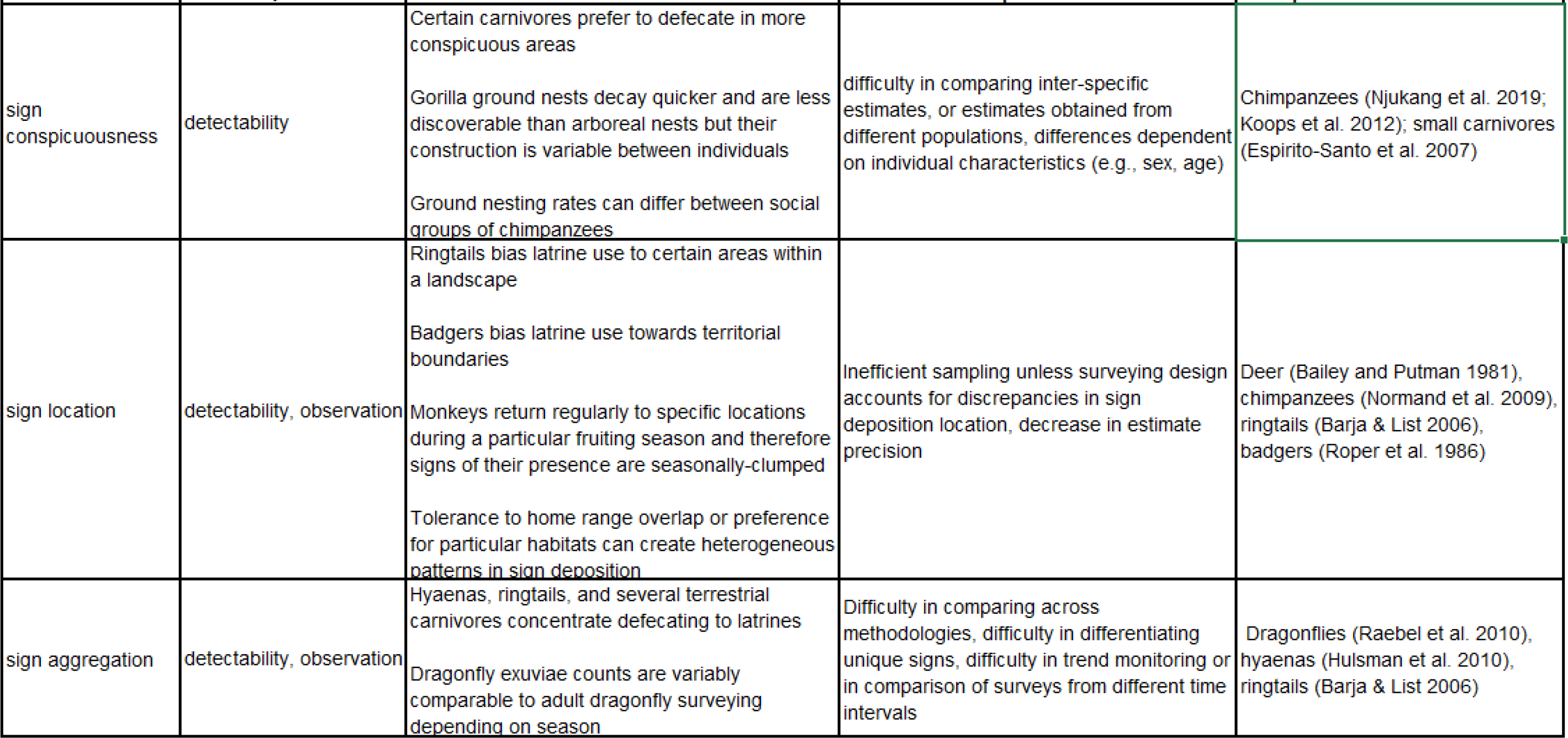
Examples of ways that behavioral variability can potentially impact the accuracy and precision of wildlife population estimates derived from sign-based monitoring, and the comparability of these estimates across contexts, the potential consequences of this variation, and measured examples of either behavioral variation and/or their impacts upon population quantification, comparison, and general estimation.

**Figure 3.**
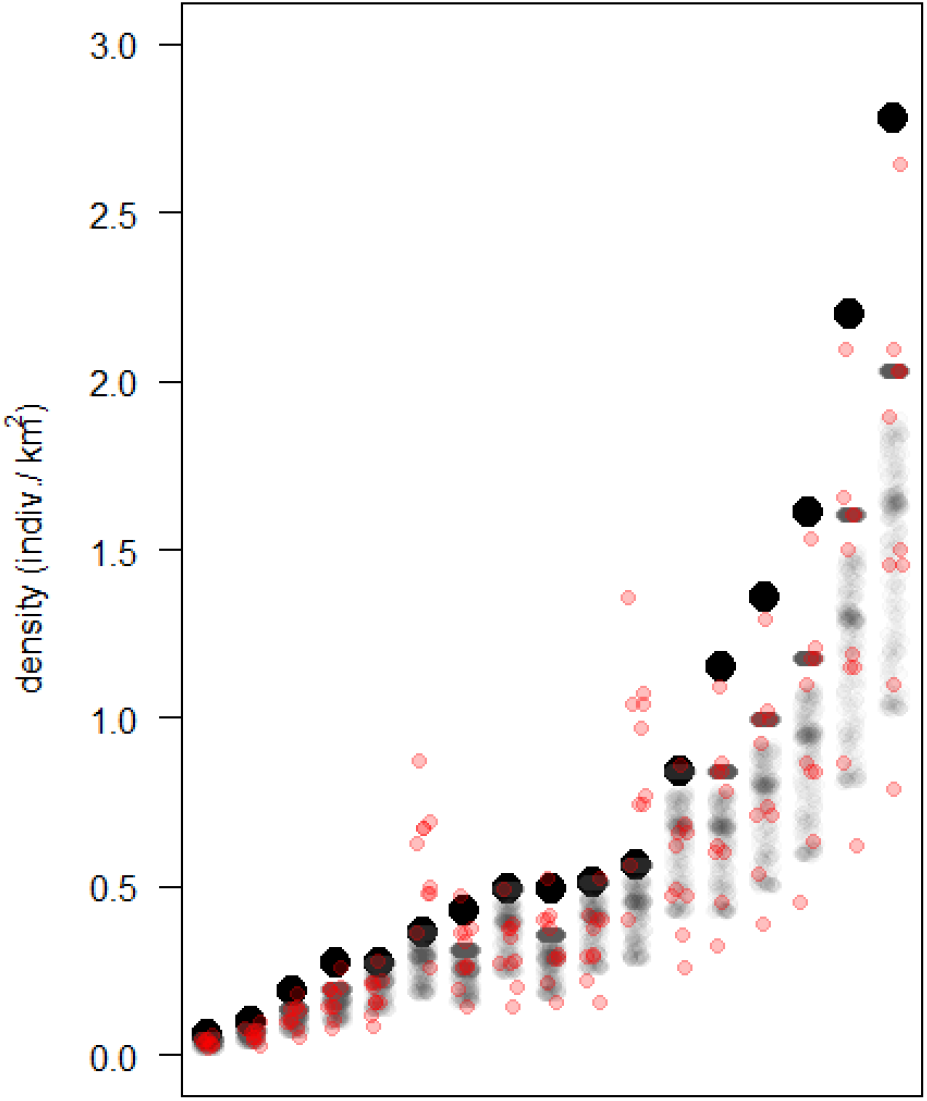
Bonobo density estimates (larger black dots, ordered low to high; data derive from density estimates provided in Table S3) and density values permuted from all observed bonobo nest construction rates (grey dots; including seasonal variation at Kokolopori; data available in Table S1 and from this study) or observed nest construction rates and nest decay rates (red dots; site-based average values only).

## DISCUSSION

When counts of animal populations depend on artefacts of behavior, wildlife monitoring methods must consider variability in animal behavior to estimate populations accurately. Our analysis demonstrates that in addition to environmental influence on sign persistence, behaviors relevant to indirect monitoring of wildlife populations can be considerably variable and significantly impact population assessment. We provide a particularly severe example, as we demonstrate that bonobos not only differ between populations in the number of nests that they construct per day, but also that within a single population this behavior is highly variable. Consequently, failure to measure and account for behavioral variability leads to potential overestimation in the number of bonobos remaining in the wild by an average of 34%, with worst-case scenarios suggesting an overestimation by up to 80%.

### The case of the bonobo

Population overestimation of this species has predominantly derived from historical reliance upon a single measure of nest production behavior being used for bonobo population density estimates. Here, we presented a second measure of nest construction rate and evaluated its seasonal variability. As the construction rate at Kokolopori was considerably higher than the single previously published estimate, and as Kokolopori does not represent any overt environmental or behavioral outlier, it is likely that we have historically overestimated bonobo abundance. While methodological comparison with camera trap studies suggests that nest counts may commonly underestimate ape densities the severity of potential overestimation described here still outpaces potential methodological underestimation from using nests as indirect signs during surveying (7.5% negative bias: Cappelle et al., 2019).

The likelihood for overestimation of bonobo populations becomes clearest when we consider the following scenarios. First, as all directly observed calculations of bonobo nest construction rates are greater than an assumed value of one nest/day, then estimates of populations using this assumed value are certainly overestimated. We found that published bonobo densities that relied on the assumption of one nest constructed per day were not only among the highest estimates, but they also were the most likely to be severely overestimated. Second, given that the Kokolopori nest construction rate is higher than the only other published rate (Mohneke & Fruth, 2008), it is likely that bonobo nest construction rates at other, non-measured sites fall closer to one of the two measured estimates than to the assumed single nest constructed per day.

### Consequences of ignoring behavioral variation in population monitoring

The example of the considerable inter- and intra-population variability in nest construction behavior has consequences for the ways in which we use population estimates derived from behavior or behavioral artefacts. Cross-population differences in indirect sign production rates considerably hamper our ability to reliably compare inter-site differences in densities. In the case of nest surveying, several authors have argued for the necessity of measuring nest decay rates locally for each survey due to environmental influence on decay (e.g., Bessone et al., 2021; Mohneke & Fruth, 2008; Plumptre & Cox, 2006), however, our results indicate that variability in nest construction behavior must likewise be considered. Without accounting for behavioral variation, our ability to discern drivers of variation in densities across populations remains obscured, which may consequently misinform conservation action.

Our results highlight further, more nuanced potential sources of biases for comparison of population estimates, which may vary from taxon to taxon. For example, we did not detect sex differences in bonobo nest construction rates, whereas this was a clear (albeit statistically untested) pattern elsewhere (Mohneke & Fruth, 2008). Inconsistencies in the influence of sex on behaviors which are also variable across populations further complicate our ability to account for these biases in population estimation, especially across populations with varying sex ratios (Plumptre & Cox, 2006).

Behavioral variability may be especially important if sign production itself can be variable across populations. For bonobos, some surveyors choose to ignore day nests during surveying because they are considered to be smaller or less robust in construction (e.g., Fruth & Hohmann, 1993; Hashimoto & Furuichi, 2002), whereas day nests at Kokolopori can appear largely indistinguishable from night nests (Supplementary Video 1; Wessling unpubl. data). Consequently, it is likely difficult to reliably differentiate day from night nests during surveying in a manner that is objective across populations. Therefore, clear (objective) decisions must be made about inclusion criteria for each indirect sign surveyed, and these decisions should reach consensus across surveys within a taxon or methodology. Further investigation into decay rate differences between day and night nests would further illuminate the impacts of decisions surrounding inclusion criteria on population monitoring.

Further, seasonality in artefact production behavior has potential impacts on comparisons of population estimates across both time and space. First, monitoring of population change relies on repeated surveying of the same population, which rests upon the assumption that variation in the observation of behavioral artifacts solely reflects variation in animal densities. However, this assumption is violated if artefact production rates vary within a site unless surveying only occurs during certain periods of the year, both within-sites (e.g., trend analysis) and between-sites. Consequently, it is necessary to understand the effect of within-population variability in relevant behaviors (e.g., nesting seasonality) on the robustness of trends derived from population resampling. In cases where behavioral variability in sign production is observed, monitoring design must, in turn, accommodate and reduce the impacts of biases introduced by this variability (e.g., by sampling during identical times of the year if nest production is seasonally-variable). The second consequence of behavioral seasonality, is that cross-site comparison of bonobo densities may be simple artefacts of differences in surveying timing. That we find seasonality in multiple artefact production behavior (ape nests: this study, defecation: Rogers 1987, Todd et al., 2008) as well as artefact decay rates (e.g., nests: Barnes, 2001; Bessone et al., 2021; Kühl et al., 2007; Nchanji Plumptre, 2001) implies that conservationists must not only account for cross-site environmental differences, but also consider potential intra-annual environmental variation during indirect survey planning, analysis, and synthesis.

Collectively, by failing to account for variation in sign production behavior we observe both clear overall biases (i.e., overestimation) in population assessment of an understudied species, as well as uncertainty in estimation and comparison of individual estimates. These effects have devastating consequences for bonobo conservation – a species for which severe data deficiencies hamper adequate conservation evaluation and prioritization (IUCN & ICCN, 2012). The demanding logistics of surveying in extremely remote regions of the Congo Basin necessitate accurate and comparative surveying because the resulting datapoints serve as the basis for extrapolation of species density across their range (e.g., Hickey et al., 2013; Nackoney & Williams, 2013). If behavioral variation in nesting imparts unaccounted-for variation in these densities, we may not only be inaccurately estimating the size of remaining populations, but also inaccurately evaluating predictors of population persistence or decline.

Our results further underline that taxa surveyed need not necessarily be behaviorally flexible for behavioral variability to be relevant for monitoring accuracy. Bonobos demonstrate muted behavioral diversity relative to chimpanzees (Hohmann & Fruth, 2003), a species well documented to exhibit a great amount of behavioral variation across its range (Kalan et al., 2020). Despite comparatively minimal ecological variation across the bonobo range (Furuichi, 2009), it is notable that bonobo nest construction rates varied considerably more between sites than the few chimpanzee construction rates recorded until now, and that bonobo nest decay rates extend across the majority of the observed variation in chimpanzee nest decay rates (Figure S1). Considering that bonobos demonstrate substantial behavioral variability with clear impacts on monitoring, and that remnants of behavior can be just as variable (e.g., decay rates) or more variable (e.g., construction rates) than behaviors in a species like the chimpanzee known for its behavioral flexibility (Kalan et al., 2020), our results underline that variability in behaviors relevant to monitoring may not necessarily follow general patterns of behavioral variability across taxa.

### How to address behavioral variability in population monitoring

While the consequences of behavioral variation on population estimation may be extreme in this example, these results should caution conservation practitioners whose methods to quantify wildlife populations may be impacted directly by wildlife behavior (e.g., temporal patterns of camera trap triggering, trappability) or which rely upon relics of behavior (e.g., dung counts). For example, circadian patterns (e.g., diurnal, cathemeral) can vary by local conditions and animals can be variably cryptic depending on environmental context (e.g., Oberosler et al., 2017; Rowcliffe et al., 2014). Collectively, these patterns point to the necessity of accounting for behavioral variation in conservation monitoring, and negate the argument that the importance of behavioral variability has a relatively minor impact among potential sources of error in indirect surveying (Marques et al., 2001; Mitchell et al., 1985).

But how can this reasonably be accomplished? It may be tempting to argue for direct sampling of relevant behaviors at each survey locality. However, the ability to directly observe wildlife behavior in a manner in which sign production rates could be calculated negates the necessity of using sign-based sampling because members of that population would be directly observable, and population estimation would therefore likely be more suitably measured using other methodologies. Unlike artefact decay rates which can be observed without needing to directly observe individual animals, collecting information on artefact production behavior does necessitate direct observation, which may only be possible in few locations. Therefore, we could consider including variance introduced by inter- and intra-site variation within the confidence intervals of computed values. However, this may not be a realistic solution, as allowing for variance of potential nest construction rates introduced 60% variance (range in density bias for ‘construction only (with seasonality)’ in Table 2) in density estimates across our sample. Expanding the confidence intervals to include this variation would render cross-sample comparison functionally meaningless, an issue that no increase in the amount of survey effort could solve (Buckland et al., 2001). Some of the uncertainty introduced by behavioral variability could be reduced via the application of multiple methodological approaches (e.g., genetic sampling, camera trapping, sign counts), to yield estimate averaging across methods, and therefore allow for cross-method validity, the avoidance of methodology-specific biases, and subsequent narrowing of possible estimate ranges (Nuñez et al. 2019). However, such an approach would require parallel surveying efforts and thus (potentially prohibitively) high monitoring costs.

Instead, a promising way forward may be to understand predictors of behavioral variation, such as environmental drivers like rainfall, which could be used as a proxy of artefact production behavior where behavioral sampling is not possible. The approach of replacing locally measured metrics of sign discoverability with environmental proxies has been previously suggested as a useful method for accommodating variability in sign decay (Meier et al. 2021; Bessone et al. 2021), and therefore may be suitably extended to proxies of behavioral variability. To accomplish this, researchers who depend on metrics of behavior in surveying should aim to increase sampling efforts of that behavior across populations where behavior can be observed, and within those populations across time periods and seasons, to characterize behavioral variation for that species. Only then, once behavioral variability can be reliably tied to predictors for a species and then modeled across time and space, could this variability be included in subsequent interpretations of inter-survey variance. Because indirect monitoring of a given species therefore must depend on estimates acquired through direct behavioral observation, long-term animal research sites must continue to be viewed as crucial components of species conservation (Campbell et al., 2011).

### Behavioral variation broadly impacts population monitoring

The impacts of behavioral variability upon population monitoring have wide reaching consequences across a variety of taxa, as all sign-based monitoring is largely dependent in some form on behavior of the individuals who leave behind these traces. A wide range of taxa are surveyed using indirect methods like tracks (e.g., ungulates: Licona et al. 2011; Reyna-Hurtado & Tanner 2007), feces/scat (e.g., elephants: Meier et al. 2021, small carnivores: Espirito-Santo et al. 2007, deer: Bailey & Putnam 1981, Marques et al. 2001, Massei & Genov 1998), and nest or drey counts (e.g., apes: Kouakou et al. 2009, this study, squirrels: Gurnell et al. 2004) that are clearly linked to behaviors that can vary. Furthermore, methodologies like hair traps (e.g., mustelids: Garcia & Mateos 2009), scent stations (e.g., bees: Almeida et al. 2019), and exuviae (Raebel et al. 2010) can also be argued to be dependent in some manner on behaviors that vary across scales (e.g., time, individual, social unit, population, species). In Table 3, we provide a few examples of the avenues through which behavioral variation can impact sign-based monitoring, with far-reaching impacts on a range of species and methodologies.

The necessity of accounting for behavioral heterogeneity across individuals has received increasing attention in the conservation literature (Kelleher et al., 2018; Merrick & Koprowski, 2017; Henrich et al. 2022). Behavioral variability can considerably impact conservation effectiveness in a number of ways, from its impact upon individual fitness, to how it affects the suitability of conservation action across contexts and populations. However, the relevance of behavioral variability has rarely been discussed in the context of conservation monitoring, with some exceptions. Behavioral variation in the form of movement, space use, and the relationship between activities that create artefacts for monitoring and landscape characteristics create considerable opportunity for biases in sign detectability, survey design, but can also increase the variance and/or decrease the accuracy of population estimates, or the fidelity of connecting animal abundances to their potential drivers (e.g., elephants: Osipova et al., 2019; alligators: Rosenblatt et al., 2013). For wildlife monitoring, issues can occur, e.g., if individuals or groups show spatiotemporal variation in territory use, variable tolerance to territory overlap, or variation in distribution of where behavioral artefacts are deposited. These patterns therefore contribute to variable stochasticity, clustering, or spatiotemporal distribution of behavioral artefacts which consequently impacts monitoring accuracy and inter-estimate comparability (Buckland et al., 2001). For example, if an ape population sleeps next to their food resources, and resources vary within the year between clumped and evenly-spaced distributions, so too would their nests. However, variation in the distribution of nests over time would also be expected to affect patterns of detectability and appropriate surveying design tailored to best monitor these signs.

Many of these examples point to a need for careful planning in surveying, quantification and accounting of potential behavioral variation and biases, and variability included in estimate precision in population monitoring. Such biases are not insurmountable if adequately acknowledged and subsequently addressed. For example, where seasonal variation in sign production behavior has been observed in a species, estimation of population densities should either be restricted to certain periods of a season or systematically averaged across a seasonal cycle. In some cases, more intensive surveying methods which may not be as susceptible to behavioral variation, such as capture-recapture methods, may offer avenues for evading the impacts of behavioral variability on population monitoring, however, such methods may not always present a feasible methodological alternative in many monitoring contexts. Moving forward, for effective and accurate indirect monitoring, it will be important that behavioral variability is considered and quantified, its impacts understood, and those impacts mitigated whenever possible, and the limitations on subsequent inference accounted for when those impacts cannot be mitigated.

Lastly, these results further underline the importance of group-level behavioral variation relative to individual-level behavior (which has been the predominant focus in the conservation literature). Whereas the impacts of individual behavioral variability upon monitoring are well documented (Biro, 2013; Carter et al., 2012; Marescot et al., 2011), the impacts of inter-population or group-level behavioral variation have additional consequences for monitoring and remain largely ignored. The relevance of group-level behavioral variation to conservation has recently gained a significant amount of attention as a potentially valuable tool to supplement traditional conservation targets if applied effectively (Brakes et al., 2019; Carvalho et al., 2022). However, our results demonstrate how group-level behavioral variation has important impacts on conservation monitoring beyond inter-individual level variation. We further illustrate the impact of temporal variation in behavior on the accuracy of population estimation. Our results support the argument that behavioral variability is relevant to conservation in other ways than just as a potential tool for advocacy or a conservation target, but has implications on our ability to effectively measure populations of concern and evaluate conservation need. Further research characterizing behavioral variation of behaviors relevant to population monitoring across individuals, time periods, populations, and environments must be performed simultaneously with measuring and mitigating its impacts upon population estimation more broadly. Given current widespread loss of wildlife, identifying how to best incorporate behavioral variation into population monitoring and conservation intervention is becoming not only pertinent but absolutely necessary.

## SUPPORTING INFORMATION

Additional information is available online in the Supporting Information section at the end of this online version. The authors are solely responsible for the content and functionality of these materials. Queries (other than absence of the material) should be directed to the corresponding author.

## Acknowledgements

This work is dedicated to the late Dr. Deborah Moore, who is deeply missed at Kokolopori. We are grateful to the Ministry of Scientific Research and Technology (MSRT) and the ICCN of the Democratic Republic of the Congo, as well as to residents of the surrounding communities for facilitating our research and grant access to the Kokolopori Bonobo Reserve. We especially thank the team of local assistants for their invaluable contribution to data collection and bonobo tracking. We kindly thank G. Tenekwetche Sop who kindly assisted with data collation for Table S2 and to Liran Samuni who coordinated data collection in the field and provided helpful feedback on earlier version of this manuscript. We sincerely thank Hjalmar Kühl, Rahel Sollmann, and two anonymous reviewers for highlighting insightful avenues of improvement to the manuscript. This work was supported by Harvard University.

## Author contributions

EW conceptualized, designed, and conducted the study; MS provided access to the data and assisted with conceptualization, and interpretation. Both authors contributed to the writing of the manuscript and have approved of the final submitted version.

## Statement on animal subjects

This study complies with the ethical standards for animal research by the standards of the American Society of Primatology and the MSRT.

## SUPPLEMENTARY MATERIAL

**Figure S1.**
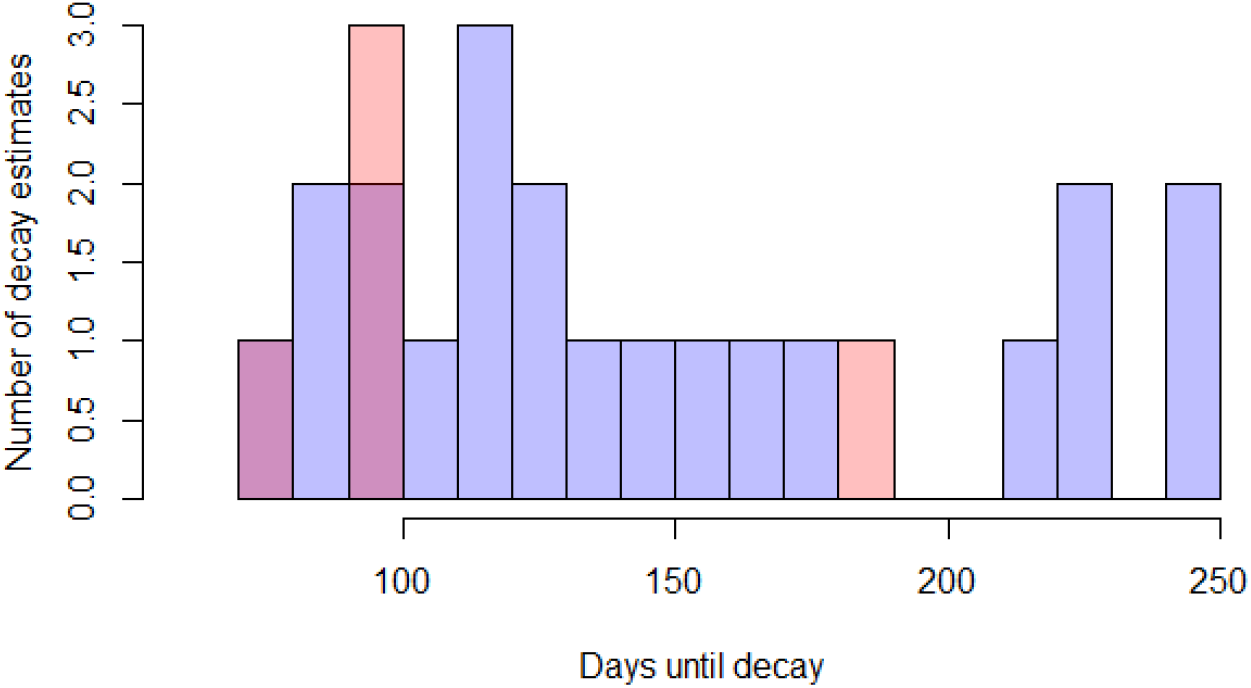
Summary of published nest decay rates of chimpanzees (blue) and bonobos (pink).

**Table S1.** Published (a) nest decay rates and (b) construction rates for chimpanzees and bonobos.

| **(a) Nest decay rates** |  |  |  |
| --- | --- | --- | --- |
| **Species** | **Days** | **Location** | **Reference** |
| Bonobo | 76 | LuiKotale, DRC | Mohneke & Fruth 2008 |
| Bonobo | 96 | LuiKotale, DRC | Bessone et al. 2021 |
| Bonobo | 99 | Lomako, DRC | van Krunkelsven 2001 |
| Bonobo | 99 | Lomako, DRC | Eriksson 1999 |
| Bonobo | 183 | Malebo, DRC | Serckx et al. 2014 |
| Chimpanzee | 73 | Taï, Côte d'Ivoire | Marchesi et al. 1995 |
| Chimpanzee | 95 | Taï, Côte d'Ivoire | Kouakou et al. 2009 |
| Chimpanzee | 161 | Taï, Côte d'Ivoire | Wild Chimpanzee Foundation; via Heinicke et al. 2019 |
| Chimpanzee | 85 | Djoroutou, Côte d'Ivoire | Wild Chimpanzee Foundation; via Heinicke et al. 2019 |
| Chimpanzee | 179 | Comoé, Côte d'Ivoire | Laupiente et al. 2020 |
| Chimpanzee | 106 | Lope, Gabon | Hall et al. 1998 |
| Chimpanzee | 114 | Belinga, Gabon | Tutin & Fernandez 1984 |
| Chimpanzee | 146 | (Goualougo) Nouabalé–Ndoki National Park, Republic of Congo | Morgan et al. 2016 |
| Chimpanzee | 90 | (Goualougo) Nouabalé–Ndoki National Park, Republic of Congo | Morgan et al. 2006 |
| Chimpanzee | 111 | Kibale, Uganda | Ghiglieri 1979; Ghiglieri 1984 |
| Chimpanzee | 243 | Dindefelo, Senegal | Heinicke et al. 2019 |
| Chimpanzee | 243/432* | Issa, Tanzania | Stewart et al. 2011 |
| Chimpanzee | 218 | Fouta Djallon, Guinea | Wild Chimpanzee Foundation; via Heinicke et al. 2019 |
| Chimpanzee | 221 | Fouta Djallon, Guinea | Ham 1998 |
| Chimpanzee | 229 | Haut Niger, Guinea | Fleury-Brugiere & Brugiere 2010 |
| Chimpanzee | 154 | Sapo, Liberia | PanAfrican Programme; via Heinicke et al. 2019 |
| Chimpanzee | 146 | Sapo, Liberia (marshes) | PanAfrican Programme; via Heinicke et al. 2019 |
| Chimpanzee | 135 | TRIDOM Landscape, Cameroon | Nzoo Dongmo et al. 2016 |
| Chimpanzee | 127 | Mbam-Djerem, Cameroon | Kamgang et al. 2020 |
| Chimpanzee | 97 | Dja, Cameroon | Bruce et al. 2018 |
| Chimpanzee | 130 | Campo Ma'an, Cameroon | Matthews & Matthews 2004 |
| Chimpanzee | 133 | Campo Ma'an, Cameroon | Nzoo Dongmo et al. 2015 |
| **(b) Nest construction rates** |  |  |  |
| **Species** | **nests/day** | **Location** | **Reference** |
| Bonobo | 1.92 | Kokolopori, DRC | *This study* |
| Bonobo | 1.37 | LuiKotale°, DRC | Mohneke & Fruth 2008 |
| Chimpanzee | 1.09 | Budongo, Uganda | Plumptre & Reynolds 1997 |
| Chimpanzee | 1.09 | Goualougo, Rep Congo | Morgan et al. 2006 |
| Chimpanzee | 1.14 | Taï, Côte d'Ivoire | Kouakou et al. 2009 |

\* Differentiated between vegetation coverage types, no site average provided; ° Previously reported as 'Lomako' in Kühl (2008).

**Table S2.** Model effect from a GLMM (Poisson error) of rainfall, group, and sex on bonobo nest construction behavior in two social groups at the Kokolopori Bonobo Reserve, DRC between 2020-2021 (12 mo.; n = 210). Statistically significant results (p ≤ 0.05) appear in bold italics.

| Predictor | Estimate ± SE | $\chi^2$ | p-value |
| --- | --- | --- | --- |
| Intercept | -1.833 ± 0.118 | - | - |
| ***Rainfall*** | 0.002 ± 0.001 | ***5.179*** | ***0.023*** |
| Group (Fekako)<sup>1</sup> | 0.061 ± 0.115 | 0.341 | 0.843 |
| Group (Kokoalongo)<sup>1</sup> | 0.041 ± 0.094 | | |
| Sex (Male)<sup>2</sup> | -0.111 ± 0.094 | 1.404 | 0.236 |

<sup>1</sup>Reference category: Ekalakala; <sup>2</sup>Reference category: Female

**Supplementary Video 1.** A bonobo constructs a day nest in the Kokolopori Bonobo Reserve, DRC. https://youtu.be/Os5sEyxQuD8

**Table S3.**
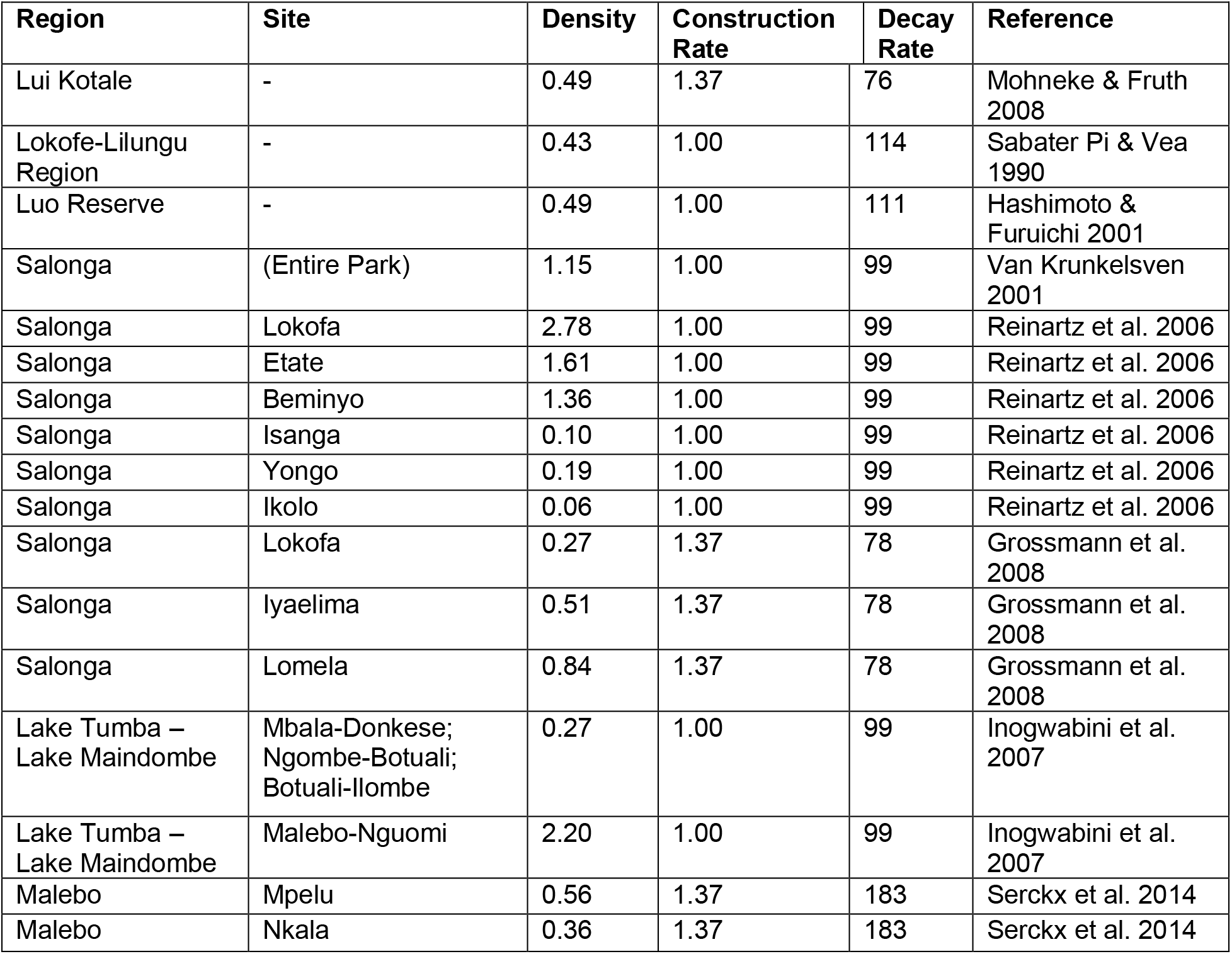
All published bonobo density estimates included in the permutations of the effects of variation in nest construction and decay rates, and the nest construction and decay rates use.

| Region | Site | Density | Construction Rate | Decay Rate | Reference |
| --- | --- | --- | --- | --- | --- |
| Lui Kotale | - | 0.49 | 1.37 | 76 | Mohneke & Fruth 2008 |
| Lokofe-Lilungu Region | - | 0.43 | 1.00 | 114 | Sabater Pi & Vea 1990 |
| Luo Reserve | - | 0.49 | 1.00 | 111 | Hashimoto & Furuichi 2001 |
| Salonga | (Entire Park) | 1.15 | 1.00 | 99 | Van Krunkelsven 2001 |
| Salonga | Lokofa | 2.78 | 1.00 | 99 | Reinartz et al. 2006 |
| Salonga | Etate | 1.61 | 1.00 | 99 | Reinartz et al. 2006 |
| Salonga | Beminyo | 1.36 | 1.00 | 99 | Reinartz et al. 2006 |
| Salonga | Isanga | 0.10 | 1.00 | 99 | Reinartz et al. 2006 |
| Salonga | Yongo | 0.19 | 1.00 | 99 | Reinartz et al. 2006 |
| Salonga | Ikolo | 0.06 | 1.00 | 99 | Reinartz et al. 2006 |
| Salonga | Lokofa | 0.27 | 1.37 | 78 | Grossmann et al. 2008 |
| Salonga | Iyaelima | 0.51 | 1.37 | 78 | Grossmann et al. 2008 |
| Salonga | Lomela | 0.84 | 1.37 | 78 | Grossmann et al. 2008 |
| Lake Tumba – Lake Maindombe | Mbala-Donkese; Ngombe-Botuali; Botuali-Ilombe | 0.27 | 1.00 | 99 | Inogwabini et al. 2007 |
| Lake Tumba – Lake Maindombe | Malebo-Nguomi | 2.20 | 1.00 | 99 | Inogwabini et al. 2007 |
| Malebo | Mpelu | 0.56 | 1.37 | 183 | Serckx et al. 2014 |
| Malebo | Nkala | 0.36 | 1.37 | 183 | Serckx et al. 2014 |

## LITERATURE CITED

Andersen, R., Hjeljord, O. Saether, B. E. (1992). Moose defecation rates in relation to habitat quality. Alces 28, 95–100.

Almeida, R. P. S., Arruda, F. V., Silva, D. P., & Coelho, B. W. T. (2019). Bees (Hymenoptera, Apoidea) in an Ecotonal Cerrado-Amazon Region in Brazil. Sociobiology, 66(3), 457–466.

Andersen, R., Hjeljord, O. & Saether, B. E. (1992). Moose defecation rates in relation to habitat quality. Alces 28, 95–100.

Baayen, R. H. (2008). Analyzing Linguistic data: A Practical Introduction to Statistics Using R. In Cambridge University Press. Cambridge.

Badji, L., Ndiaye, P. I., Lindshield, S. M., Ba, C. T., & Pruetz, J. D. (2018). Savanna chimpanzee (Pan troglodytes verus) nesting ecology at Bagnomba (Kedougou, Senegal). Primates, 59(3), 235–241.

Bailey, R. E., & Putman, R. J. (1981). Estimation of fallow deer (Dama dama) populations from faecal accumulation. Journal of Applied Ecology, 697–702.

Barnes, R. F. (2001). How reliable are dung counts for estimating elephant numbers? African Journal of Ecology, 39(1), 1–9.

Barocas, A., & Ben-David, M. (2021). Social Structure of Marine Otters: Inter and Intraspecific Variation. In: Ethology and Behavioral Ecology of Sea Otters and Polar Bears. Springer, Cham, pp. 83–105.

Berger, J. (1978). Group size, foraging, and antipredator ploys: an analysis of bighorn sheep decisions. Behavioral Ecology and Sociobiology, 91–99.

Bersacola, E., Hill, C. M., Nijman, V., & Hockings, K. J. (2022). Examining primate community occurrence patterns in agroforest landscapes using arboreal and terrestrial camera traps. Landscape Ecology, 1–19.

Bessone, M., Booto, L., Santos, A. R., Kühl, H. S., & Fruth, B. (2021). No time to rest: How the effects of climate change on nest decay threaten the conservation of apes in the wild. PLoS One, 16(6), e0252527.

Biro, P. A. (2013). Are most samples of animals systematically biased? Consistent individual trait differences bias samples despite random sampling. Oecologia, 171(2), 339–345.

Bogart, S. L., & Pruetz, J. D. (2011). Insectivory of savanna chimpanzees (Pan troglodytes verus) at Fongoli, Senegal. American Journal of Physical Anthropology, 145(1), 11–20.

Bonesi, L., & Macdonald, D. W. (2004). Evaluation of sign surveys as a way to estimate the relative abundance of American mink (Mustela vison). Journal of Zoology, 262(1), 65–72.

Brakes, P., Dall, S. R., Aplin, L. M., Bearhop, S., Carroll, E. L., Ciucci, P., Fishlock, V., Ford, J. K., Garland, E. C., & Keith, S. A. (2019). Animal cultures matter for conservation. Science, 363(6431), 1032–1034.

Buckland, S. T., Anderson, D. R., Burnham, K. P., Laake, J. L., Borchers, D. L., & Thomas, L. (2001). Introduction to distance sampling: estimating abundance of biological populations. Oxford: Oxford University Press.

Buckland, S. T., Rexstad, E. A., Marques, T. A., & Oedekoven, C. S. (2015). Distance sampling: methods and applications (Vol. 431): Springer.

Campbell, G., Kühl, H., Diarrassouba, A., N’Goran, P K., & Boesch, C. (2011). Long-term research sites as refugia for threatened and over-harvested species. Biology Letters, 7(5), 723–726.

Campos-Candela, A., Palmer, M., Balle, S., & Alós, J. (2018). A camera-based method for estimating absolute density in animals displaying home range behaviour. Journal of Animal Ecology, 87(3), 825–837.

Cappelle, N., Després-Einspenner, M. L., Howe, E. J., Boesch, C., & Kühl, H. S. (2019). Validating camera trap distance sampling for chimpanzees. American Journal of Primatology, 81(3), e22962.

Carter, A. J., Heinsohn, R., Goldizen, A. W., & Biro, P. A. (2012). Boldness, trappability and sampling bias in wild lizards. Animal Behaviour, 83, 1051e1058.

Carvalho, S., Wessling, E., Abwe, E. E., Almeida-Warren, K., Arandjelovic, M., Boesch, C., Danquah, E., Diallo, M. S., Hobaiter, C., & Hockings, K., Humle, T., Ashegbofe Ikemeh, R., Kalan, A. K., Luncz, L., Ohashi, G., Pascual-Garrido, A., Piel, A., Samuni, L., Soiret, S., Sanz, C. K. Koops. (2022). Using non-human culture in conservation requires careful and concerted action. Conservation Letters.

Cochran, G. A., & Stains, H. J. (1961). Deposition and decomposition of fecal pellets by cottontails. The Journal of Wildlife Management, 25(4), 432–435.

Crowcroft, P., & Jeffers, J. N. (1961, December). Variability in the behaviour of wild house mice (Mus musculus L.) towards live traps. In Proceedings of the Zoological Society of London (Vol. 137, No. 4, pp. 573–582). Oxford, UK: Blackwell Publishing Ltd.

Crunchant, A. S., Borchers, D., Kühl, H., & Piel, A. (2020). Listening and watching: do camera traps or acoustic sensors more efficiently detect wild chimpanzees in an open habitat? Methods in Ecology and Evolution, 11(4), 542–552.

Deutsch, S., Pearson, H., & Würsig, B. (2014). Development of leaps in dusky dolphin (Lagenorhynchus obscurus) calves. Behaviour, 151(11), 1555–1577.

Dobson, A. J. (2002). An Introduction to Generalized Linear Models. Boca Raton, FL: Chapman & Hall/CRC.

Doming, J., & Harris, S. (2019). Individual and seasonal variation in contact rate, connectivity and centrality in red fox (Vulpes vulpes) social groups. Scientific Reports, 9(1), 1–11.

Espírito-Santo, C., Rosalino, L. M., & Santos-Reis, M. (2007). Factors affecting the placement of common genet latrine sites in a Mediterranean landscape in Portugal. Journal of mammalogy, 88(1), 201–207.

Fruth, B., & Hohmann, G. (1993). Ecological and behavioral aspects of nest building in wild bonobos (Pan paniscus). Ethology, 94(2), 113–126.

Fruth, B., Hickey, J., André, C., Furuichi, T., Hart, J., Hart, T., Kühl, H., Maisels, F., Nackoney, J., & Reinartz, G. (2016). Pan paniscus. IUCN Red List.

Furuichi, T. (2009). Factors underlying party size differences between chimpanzees and bonobos: a review and hypotheses for future study. Primates, 50(3), 197–209.

García, P., & Mateos, I. (2009). Evaluation of three indirect methods for surveying the distribution of the least weasel Mustela nivalis in a Mediterranean area. Small Carnivore Conservation, 40, 22–26.

Gurnell, J., Lurz, P. W. W., Shirley, M. D. F., Cartmel, S., Garson, P. J., Magris, L., & Steele, J. (2004). Monitoring red squirrels Sciurus vulgaris and grey squirrels Sciurus carolinensis in Britain. Mammal Review, 34(1-2), 51–74.

Hammond, T. T., Curtis, M. J., Jacobs, L. E., Tobler, M. W., Swaisgood, R. R., & Shier, D. M. (2021). Behavior and detection method influence detection probability of a translocated, endangered amphibian. Animal Conservation, 24(3), 401–411.

Hashimoto, C., & Furuichi, T. (2002). Current situation of bonobos in the Luo reserve, Equateur, democratic republic of Congo. In All Apes Great and Small (pp. 83–90): Springer.

Hayward, M. W. & Marlow, N. (2014) Will dingoes really conserve wildlife and can our methods tell? Journal of Applied Ecology, 51, 835–838.

Hedges, S., Tyson, M. J., Sitompul, A. F., Kinnaird, M. F., & Gunaryadi, D. (2005). Distribution, status, and conservation needs of Asian elephants (Elephas maximus) in Lampung Province, Sumatra, Indonesia. Biological Conservation, 124(1), 35–48.

Henrich, M., Hartig, F., Dormann, C. F., Kühl, H. S., Peters, W., Franke, F., … & Heurich, M. D. (2022). Deer Behavior Affects Density Estimates With Camera Traps, but Is Outwighted by Spatial Variability. Frontiers in Ecology and Evolution.

Hickey, J. R., Nackoney, J., Nibbelink, N. P., Blake, S., Bonyenge, A., Coxe, S., Dupain, J., Emetshu, M., Furuichi, T., & Grossmann, F. (2013). Human proximity and habitat fragmentation are key drivers of the rangewide bonobo distribution. Biodiversity and Conservation, 22(13), 3085–3104.

Hohmann, G., & Fruth, B. (2003). Culture in bonobos? Between-species and within-species variation in behavior. Current Anthropology, 44(4), 563–571.

Howe, E. J., Buckland, S. T., Després-Einspenner, M. L., & Kühl, H. S. (2017). Distance sampling with camera traps. Methods in Ecology and Evolution, 8(11), 1558–1565.

Hulsman, A., Dalerum, F., Swanepoel, L., Ganswindt, A., Sutherland, C., & Paris, M. (2010). Patterns of scat deposition by brown hyaenas Hyaena brunnea in a mountain savannah region of South Africa. Wildlife Biology, 16(4), 445–451.

Inogwabini, B.-I., Bewa, M., Longwango, M., Abokome, M., & Vuvu, M. (2008). The bonobos of the Lake Tumba–Lake Maindombe hinterland: threats and opportunities for population conservation. In The Bonobos (pp. 273–290): Springer.

IUCN, & ICCN. (2012). Bonobo (Pan paniscus): conservation strategy 2012-2022. IUCN/ICCN, Gland.

Kalan, A. K., Hohmann, G., Arandjelovic, M., Boesch, C., McCarthy, M. S., Agbor, A., … & Kühl, H. S. (2019). Novelty response of wild African apes to camera traps. Current Biology, 29(7), 1211–1217.

Kalan, A. K., Kulik, L., Arandjelovic, M., Boesch, C., Haas, F., Dieguez, P., Barratt, C. D., Abwe, E. E., Agbor, A., Angedakin, S., Aubert, F., Ayimisin, E. A., Bailey, E., Bessone, M., Brazzola, G., Buh, V. E., Chancellor, R., Cohen, H., Coupland, C., Curran, B., et al. (2020). Environmental variability supports chimpanzee behavioural diversity. Nature Communications, 11(1), 1–10.

Kamgang, S. A., Carme, T. C., Bobo, K. S., Abwe, E. E., Gonder, M. K., & Sinsin, B. (2020). Assessment of in situ nest decay rate for chimpanzees (Pan troglodytes ellioti Matschie, 1914) in Mbam-Djerem National Park, Cameroon: implications for long-term monitoring. Primates, 61(2), 189–200.

Kelleher, S. R., Silla, A. J., & Byrne, P. G. (2018). Animal personality and behavioral syndromes in amphibians: a review of the evidence, experimental approaches, and implications for conservation. Behavioral Ecology and Sociobiology, 72(5), 1–26.

Koops, K., McGrew, W. C., Matsuzawa, T., & Knapp, L. A. (2012). Terrestrial nest-building by wild chimpanzees (Pan troglodytes): implications for the tree-to-ground sleep transition in early hominins. American Journal of Physical Anthropology, 148(3), 351–361.

Kouakou, C. Y., Boesch, C., & Kühl, H. (2009). Estimating chimpanzee population size with nest counts: validating methods in Taï National Park. American Journal of Primatology, 71(6), 447–457.

Kühl, H. (2008). Best practice guidelines for the surveys and monitoring of great ape populations: IUCN.

Kühl, H. S., Todd, A., Boesch, C., & Walsh, P. D. (2007). Manipulating decay time for efficient large-mammal density estimation: gorillas and dung height. Ecological Applications, 17(8), 2403–2414.

Laing, S., Buckland, S., Burn, R., Lambie, D., & Amphlett, A. (2003). Dung and nest surveys: estimating decay rates. Journal of Applied Ecology, 40(6), 1102–1111.

Lea, J. S., Wetherbee, B. M., Queiroz, N., Burnie, N., Aming, C., Sousa, L. L., … & Shivji, M. S. (2015). Repeated, long-distance migrations by a philopatric predator targeting highly contrasting ecosystems. Scientific Reports, 5(1), 1–11.

Licona, M., McCleery, R., Collier, B., Brightsmith, D. J., & Lopez, R. (2011). Using ungulate occurrence to evaluate community-based conservation within a biosphere reserve model. Animal Conservation, 14(2), 206–214.

Luccas, V., & Izar, P. (2021). Black capuchin monkeys dynamically adjust group spread throughout the day. Primates, 62(5), 789–799.

Marescot, L., Pradel, R., Duchamp, C., Cubaynes, S., Marboutin, E., Choquet, R., Miquel, C., & Gimenez, O. (2011). Capture–recapture population growth rate as a robust tool against detection heterogeneity for population management. Ecological Applications, 21(8), 2898–2907.

Marques, F. F., Buckland, S. T., Goffin, D., Dixon, C. E., Borchers, D. L., Mayle, B. A., & Peace, A. J. (2001). Estimating deer abundance from line transect surveys of dung: sika deer in southern Scotland. Journal of Applied Ecology, 349–363.

Massei, G., & Genov, P. V. (1998). Fallow deer (Dama dama) winter defecation rate in a Mediterranean area. Journal of Zoology, 245(2), 209–211.

Mayle, A. B., Doney, J., Lazarus, G., Peace, A. J. & Smith, D. E. (1996). Fallow deer (Dama dama) defecation rate and its use in determining population size. Suppl. Ric. Biol. Selvaggina 25, 63–78.

Meier, A. C., Shirley, M. H., Beirne, C., Breuer, T., Lewis, M., Masseloux, J., … & Poulsen, J. R. (2021). Improving population estimates of difficult-to-observe species: A dung decay model for forest elephants with remotely sensed imagery. Animal Conservation, 24(6), 1032–1045.

Merrick, M. J., & Koprowski, J. L. (2017). Should we consider individual behavior differences in applied wildlife conservation studies? Biological Conservation, 209, 34–44.

Mitchell, B. D., Rowe, J. J., Ratcliffe, P. R. & Hinge, M. (1985). Defecation frequency in roe deer (Capreolus capreolus) in relation to the accumulation rates of faecal deposits. Journal of Zoology, 207, 1–7.

Moehlman, P. D. (1998). Feral asses (Equus africanus): intraspecific variation in social organization in arid and mesic habitats. Applied Animal Behaviour Science, 60(2-3), 171–195.

Moeller, A. K., Lukacs, P. M., & Horne, J. S. (2018). Three novel methods to estimate abundance of unmarked animals using remote cameras. Ecosphere, 9(8), e02331.

Mohneke, M., & Fruth, B. (2008). Bonobo (Pan paniscus) density estimation in the SW-Salonga National Park, Democratic Republic of Congo: common methodology revisited. In The Bonobos (pp. 151–166): Springer.

Morgan, D., Sanz, C., Onononga, J. R., & Strindberg, S. (2006). Ape abundance and habitat use in the Goualougo Triangle, Republic of Congo. International Journal of Primatology, 27(1), 147–179.

Morgan, D., Sanz, C., Onononga, J. R., & Strindberg, S. (2016). Factors influencing the survival of sympatric gorilla (Gorilla gorilla gorilla) and chimpanzee (Pan troglodytes troglodytes) nests. International Journal of Primatology, 37(6), 718–737.

Nackoney, J., & Williams, D. (2013). A comparison of scenarios for rural development planning and conservation in the Democratic Republic of the Congo. Biological Conservation, 164, 140–149.

Nakashima, Y., Fukasawa, K., & Samejima, H. (2018). Estimating animal density without individual recognition using information derivable exclusively from camera traps. Journal of Applied Ecology, 55(2), 735–744.

Nchanji, A. C., & Plumptre, A. J. (2001). Seasonality in elephant dung decay and implications for censusing and population monitoring in south-western Cameroon. African Journal of Ecology, 39(1), 24–32.

Njukang, A. P., Angwafor, T. E., Richard, S. E. I. N. O., Akwanjoh, A. K. L., & Chuo, M. D. (2019). Effects of anthropogenic activities on chimpanzee nest location in the Tofala hill wildlife sanctuary (THWS), South West Region, Cameroon. International Journal of Forest, Animal and Fisheries Research, 3(1), 1–9.

Normand, E., Ban, S. D., & Boesch, C. (2009). Forest chimpanzees (Pan troglodytes verus) remember the location of numerous fruit trees. Animal Cognition, 12(6), 797–807.

Nuñez, C. L., Froese, G., Meier, A. C., Beirne, C., Depenthal, J., Kim, S., … & Poulsen, J. R. (2019). Stronger together: comparing and integrating camera trap, visual, and dung survey data in tropical forest communities. Ecosphere, 10(12), e02965.

Oberosler, V., Groff, C., Iemma, A., Pedrini, P., & Rovero, F. (2017). The influence of human disturbance on occupancy and activity patterns of mammals in the Italian Alps from systematic camera trapping. Mammalian Biology, 87(1), 50–61.

Osipova, L., Okello, M., Njumbi, S., Ngene, S., Western, D., Hayward, M., & Balkenhol, N. (2019). Using step-selection functions to model landscape connectivity for African elephants: accounting for variability across individuals and seasons. Animal Conservation, 22(1), 35–48.

Plumptre, A. J. (2000). Monitoring mammal populations with line transect techniques in African forests. Journal of Applied Ecology, 37(2), 356–368.

Plumptre, A. J., & Cox, D. (2006). Counting primates for conservation: primate surveys in Uganda. Primates, 47(1), 65–73.

Poggenburg, C., Nopp-Mayr, U., Coppes, J., & Sachser, F. (2018). Shit happens … and persists: decay dynamics of capercaillie (Tetrao urogallus L.) droppings under natural and artificial conditions. European Journal of Wildlife Research, 64(3), 1–16.

Raebel, E. M., Merckx, T., Riordan, P., Macdonald, D. W., & Thompson, D. J. (2010). The dragonfly delusion: why it is essential to sample exuviae to avoid biased surveys. Journal of Insect Conservation, 14(5), 523–533.

Reinartz, G. E., Isia, I. B., Ngamankosi, M., & Wema, L. W. (2006). Effects of forest type and human presence on bonobo (Pan paniscus) density in the Salonga National Park. International Journal of Primatology, 27(2), 603–634.

Reyna-Hurtado, R., & Tanner, G. W. (2007). Ungulate relative abundance in hunted and non-hunted sites in Calakmul Forest (Southern Mexico). Biodiversity and Conservation, 16(3), 743–756.

Rogers, L. L. (1987). Seasonal changes in defecation rates of free-ranging white-tailed deer. J. Wildl. Manage. 51, 330–333.

Roper, T. J., Shepherdson, D. J., & Davies, J. M. (1986). Scent marking with faeces and anal secretion in the European badger (Meles meles): seasonal and spatial characteristics of latrine use in relation to territoriality. Behaviour, 94–117.

Rosenblatt, A. E., Heithaus, M. R., Mazzotti, F. J., Cherkiss, M., & Jeffery, B. M. (2013). Intra-population variation in activity ranges, diel patterns, movement rates, and habitat use of American alligators in a subtropical estuary. Estuarine, Coastal and Shelf Science, 135, 182–190.

Rowcliffe, J. M., Kays, R., Kranstauber, B., Carbone, C., & Jansen, P. A. (2014). Quantifying levels of animal activity using camera trap data. Methods in Ecology and Evolution, 5(11), 1170–1179.

Samuni, L., Wegdell, F., & Surbeck, M. (2020). Behavioural diversity of bonobo prey preference as a potential cultural trait. eLife, 9, e59191.

Schmaljohann, H., Liechti, F., Bächler, E., Steuri, T., & Bruderer, B. (2008). Quantification of bird migration by radar–a detection probability problem. Ibis, 150(2), 342–355.

Serckx, A., Huynen, M.-C., Bastin, J.-F., Hambuckers, A., Beudels-Jamar, R. C., Vimond, M., Raynaud, E., & Kühl, H. S. (2014). Nest grouping patterns of bonobos (Pan paniscus) in relation to fruit availability in a forest-savannah mosaic. PLoS One, 9(4), e93742.

Stewart, F. A., Piel, A. K., Azkarate, J. C., & Pruetz, J. D. (2018). Savanna chimpanzees adjust sleeping nest architecture in response to local weather conditions. American Journal of Physical Anthropology, 166(3), 549–562.

Stolwijk, A., Straatman, H., & Zielhuis, G. (1999). Studying seasonality by using sine and cosine functions in regression analysis. Journal of Epidemiology & Community Health, 53(4), 235–238.

Strindberg, S., Maisels, F., Williamson, E. A., Blake, S., Stokes, E. J., Aba’a, R., Abitsi, G., Agbor, A., Ambahe, R. D., & Bakabana, P. C. (2018). Guns, germs, and trees determine density and distribution of gorillas and chimpanzees in Western Equatorial Africa. Science Advances, 4(4), eaar2964.

Surbeck, M., Coxe, S., & Lokasola, A. L. (2017). Lonoa: The Establishment of a Permanent Field Site for Behavioural Research on Bonobos in the Kokolopori Bonobo Reserve. Pan Africa News, 24(2), 13–15.

Todd, A. F., Kühl, H. S., Cipolletta, C., & Walsh, P. D. (2008). Using dung to estimate gorilla density: modeling dung production rate. International Journal of Primatology, 29(2), 549–563.

Van Krunkelsven, E. (2001). Density estimation of bonobos (Pan paniscus) in Salonga National Park, Congo. Biological Conservation, 99(3), 387–391.

Viquerat, S. M. A., Muller, M., Kiffner, C., Waltert, M., & Bobo, K. S. (2012). Estimating forest duiker (Cephalophinae) density in Korup National Park: a case study on the performance of three line transect methods. South African Journal of Wildlife Research, 42(1), 1–10.

Walsh, P. D., & White, L. J. (2005). Evaluating the steady state assumption: simulations of gorilla nest decay. Ecological Applications, 15(4), 1342–1350.

Wong, S. T., Belant, J. L., Sollmann, R., Mohamed, A., Niedballa, J., Mathai, J., … & Wilting, A. (2019). Influence of body mass, sociality, and movement behavior on improved detection probabilities when using a second camera trap. Global Ecology and Conservation, 20, e00791.

## LITERATURE CITED

Bates, D., Mächler, M., Bolker, B., & Walker, S. (2015). Fitting Linear Mixed-Effects Models Using lme4. Journal of Statistical Software, 67(i01).

Bruce, T., Ndjassi, C., Fowler, A., Ndimbe, M., Fankem, O., Mbodbda, R. B. T., Kobla, A.-S., Puemo, F. A. W., Lushimba, A., Amin, R., Wacher, T., Grange-Chamfray, S., & Olson, D. (2018). Faunal Inventory of the Dja Faunal Reserve, Cameroon – 2018. Ministry of Forests and Wildlife (MINFOF), Zoological Society of London – Cameroon Country Programme, African Wildlife Foundation, Yaoundé, Cameroon.

Dobson, A. J., & Barnett, A. G. (2018). An introduction to generalized linear models: CRC press.

Eriksson, J. (1999). A survey of the forest and census of the bonobo (Pan paniscus) population between the Lomako and Yekokora Rivers in the Equateur Province, DR Congo. MS thesis, Univ. of Uppsala.

Fleury-Brugiere, M.-C., & Brugiere, D. (2010). High population density of Pan troglodytes verus in the Haut Niger National Park, Republic of Guinea: implications for local and regional conservation. International Journal of Primatology, 31(3), 383–392.

Ghiglieri, M. P. (1984). 6. Feeding Ecology and Sociality of Chimpanzees in Kibale Forest, Uganda. In Adaptations for foraging in nonhuman primates (pp. 161–194): Columbia University Press.

Grossmann, F., Hart, J. A., Vosper, A., & Ilambu, O. (2008). Range occupation and population estimates of bonobos in the Salonga National Park: application to large-scale surveys of bonobos in the Democratic Republic of Congo. In The bonobos (pp. 189–216). Springer, New York, NY.

Hall, J. S., White, L. J., Inogwabini, B.-I., Omari, I., Morland, H. S., Williamson, E. A., Saltonstall, K., Walsh, P., Sikubwabo, C., & Bonny, D. (1998). Survey of Grauer’s gorillas (Gorilla gorilla graueri) and eastern chimpanzees (Pan troglodytes schweinfurthi) in the Kahuzi-Biega National Park lowland sector and adjacent forest in eastern Democratic Republic of Congo. International Journal of Primatology, 19(2), 207–235.

Ham, R. (1998). Chimpanzee survey in the Republic of Guinea. Report for the European Union.

Heinicke, S., Mundry, R., Boesch, C., Amarasekaran, B., Barrie, A., Brncic, T., Brugière, D., Campbell, G., Carvalho, J., & Danquah, E. (2019). Characteristics of positive deviants in western chimpanzee populations. Frontiers in Ecology and Evolution, 7, 16.

Inogwabini, B. I., Matungila, B., Mbende, L., Abokome, M., & wa Tshimanga, T. (2007). Great apes in the Lake Tumba landscape, Democratic Republic of Congo: newly described populations. Oryx, 41(4), 532–538.

Kouakou, C. Y., Boesch, C., & Kuehl, H. (2009). Estimating chimpanzee population size with nest counts: validating methods in Taï National Park. American Journal of Primatology: Official Journal of the American Society of Primatologists, 71(6), 447–457.

Kühl, H. (2008). Best practice guidelines for the surveys and monitoring of great ape populations. IUCN.

Lapuente, J., Ouattara, A., Köster, P. C., & Linsenmair, K. E. (2020). Status and distribution of Comoé Chimpanzees: combined use of transects and camera traps to quantify a low-density population in savanna-forest mosaic. Primates, 61(5), 647–659.

Marchesi, P., Marchesi, N., Fruth, B., & Boesch, C. (1995). Census and distribution of chimpanzees in Cote d’Ivoire. Primates, 36(4), 591–607.

Matthews, A., & Matthews, A. (2004). Survey of gorillas (Gorilla gorilla gorilla) and chimpanzees (Pan troglodytes troglodytes) in Southwestern Cameroon. Primates, 45(1), 15–24.

Nzooh Dongmo, Z. L., N’goran, K. P., Etoga, G., Belinga, J. P., Fouda, E., Bandjouma, M., & Dongmo, P. (2016). Les populations de grands et moyens mammifères dans le segment Cameroun du Paysage TRIDOM: Forêt de Ngoyla-Mintom, PN Boumba Bek et PN Nki et leurs zones périphériques. Rapport Technique. Yaoundé, Cameroun.

Nzooh Dongmo, Z. L., N’Goran, P. K., Fondja, C., & Nkono, J. (2015). Evaluation de la Dynamique des Populations de Grands et Moyens Mammiferes dans le Domaine Dorestier Permanent de l’Unite Technique Operationnelle Campo Ma’an. Service de Conservation du Parc National de Campo Ma’an, WWF Regional Office for Africa. Yaounde, Cameroon.

Plumptre, A. J., & Reynolds, V. (1997). Nesting behavior of chimpanzees: implications for censuses. International Journal of Primatology, 18(4), 475–485.

Reinartz, G. E., Isia, I. B., Ngamankosi, M., & Wema, L. W. (2006). Effects of forest type and human presence on bonobo (Pan paniscus) density in the Salonga National Park1. International Journal of Primatology, 27(2), 603–634.

Sabater-Pi, J., & Vea, J. (1990). Nest building and population estimates of the bonobo from Lokofe Lilungu-Ikomaloki region of Zaire. Primate Conservation, 11, 43–48.

Serckx, A., Huynen, M. C., Bastin, J. F., Hambuckers, A., Beudels-Jamar, R. C., Vimond, M., … & Kühl, H. S. (2014). Nest grouping patterns of bonobos (Pan paniscus) in relation to fruit availability in a forest-savannah mosaic. PloS one, 9(4), e93742.

Stewart, F. A., Piel, A. K., & McGrew, W. C. (2011). Living archaeology: artefacts of specific nest site fidelity in wild chimpanzees. Journal of Human Evolution, 61(4), 388–395.

Tutin, C. E., & Fernandez, M. (1984). Nationwide census of gorilla (Gorilla g. gorilla) and chimpanzee (Pan t. troglodytes) populations in Gabon. American Journal of Primatology, 6(4), 313–336.

